# A Multi-Layered Atlas of Spatial Regulatory Programs and Therapeutic Vulnerabilities in Glioblastoma

**DOI:** 10.1101/2025.09.27.674707

**Authors:** Linan Zhang, Matthew Lu, Xiaojun Ma, Hatice Osmanbeyoglu

## Abstract

Glioblastoma (GBM) is marked by profound spatial heterogeneity. This complexity fuels tumor progression and therapy resistance. Although recent spatial transcriptomics (ST) studies have defined transcriptional niches, or metaprograms, the systems level regulatory logic linking upstream signaling, transcriptional programs, and therapeutic vulnerabilities remains unresolved. Here, we introduce an integrated, multi layered framework that unifies transcription factor and pathway activity inference (STAN and SPAN) with ligand/receptor and drug target mapping, thereby constructing a systems-level regulatory atlas of GBM across 26 tumors. This atlas uncovers spatial niches organized around transcriptional hubs and signaling programs, including hypoxic mesenchymal regions coordinated by HIF1A and SOX2 with mTORC1 signaling, macrophage rich areas driven by STAT3 mediated immune modulation, and astrocyte-like states shaped by lipid metabolic regulators such as SREBF2. By coupling this regulatory atlas to drug target predictions, we identify niche specific therapeutic vulnerabilities, such as VEGF blockade in HIF1A high regions and MAO-B inhibition in mesenchymal states. Finally, we provide an open, interactive web resource to make these data broadly accessible. This work provides a generalizable blueprint for linking spatial regulation to therapeutic hypotheses across diverse human diseases.

## Introduction

Glioblastoma (GBM) is the most aggressive primary brain malignancy, comprising 48% of all cases and causing over 10,000 deaths annually in the United States^1,2^ despite multimodal therapy. A major contributor to this poor outcome is the intra-tumoral heterogeneity, which includes not only malignant cells but also a dynamic tumor microenvironment (TME) of immune, stromal, and glial cells^3-7^. These diverse components engage in spatially organized interactions that fuel tumor progression, therapy resistance, and immune evasion. Overcoming this complexity requires therapeutic strategies that simultaneously address molecular drivers and TME organization.

Spatial transcriptomics (ST) has recently emerged as a powerful technology to map gene expression within intact tissue architecture, enabling systems-level interrogation of GBM’s spatial complexity^8,9^. This approach has revealed robust gene expression signatures, or *metaprograms* (MPs)^8^, that capture malignant and non-malignant states within the TME. While these studies have successfully defined transcriptional niches, they have not resolved the systems-level regulatory logic that connects upstream signaling to TF-driven programs and ultimately to therapeutic vulnerabilities.

We previously developed a computational framework called STAN (Spatially informed Transcription Factor Activity Network) to predict spot-specific TF activities from ST data^10^. Here, we extend this approach by introducing SPAN (Spatially informed Pathway Activity Network) to infer spatially localized pathway activities. Then, we apply this integrated framework to 26 GBM samples to build a multi-layered regulatory atlas, link regulatory map to therapeutic vulnerabilities^11^, and provide an open resource for the community^11^ (**Figure 1**). Beyond GBM, this integrated framework establishes a generalizable systems-level strategy for connecting spatially resolved regulatory programs to experimentally testable therapeutic hypotheses across diverse diseases.

**Figure 1.**
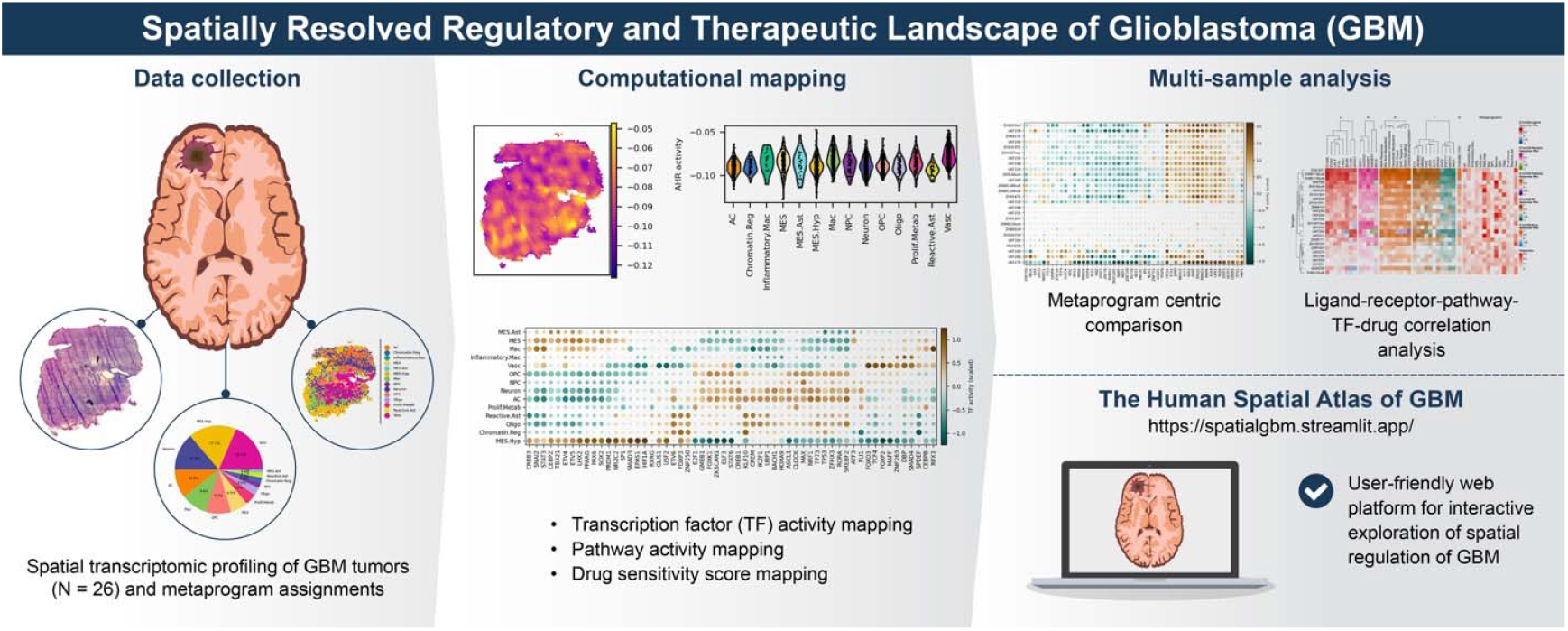
Multi-layered framework for mapping spatial regulation in glioblastoma. Schematic of the integrative pipeline combining transcription factor activity inference (STAN), pathway activity inference (SPAN), ligand–receptor mapping, and drug–target predictions. The framework links spatial gene expression and metaprograms to regulatory networks and therapeutic vulnerabilities, producing a systems-level atlas of GBM tumor ecosystems.

## Results

### Regulatory Programs of GBM TME

We analyzed Visium spatial transcriptomics (ST; 10x Genomics) samples (n = 26) from two independent GBM cohorts^8,9^. To define spatially resolved cellular states, we annotated spots using metaprogram (MP) assignments from Greenwald et al.^8^ (**Methods, Figure 2A**). MPs are recurrent transcriptional programs identified across tumors via non-negative matrix factorization, representing modules of co-expressed genes that capture both malignant and non-malignant states within the GBM TME. By providing a structured framework for spot annotation, MPs enable systematic comparisons of cellular states across spatial contexts.

**Figure 2.**
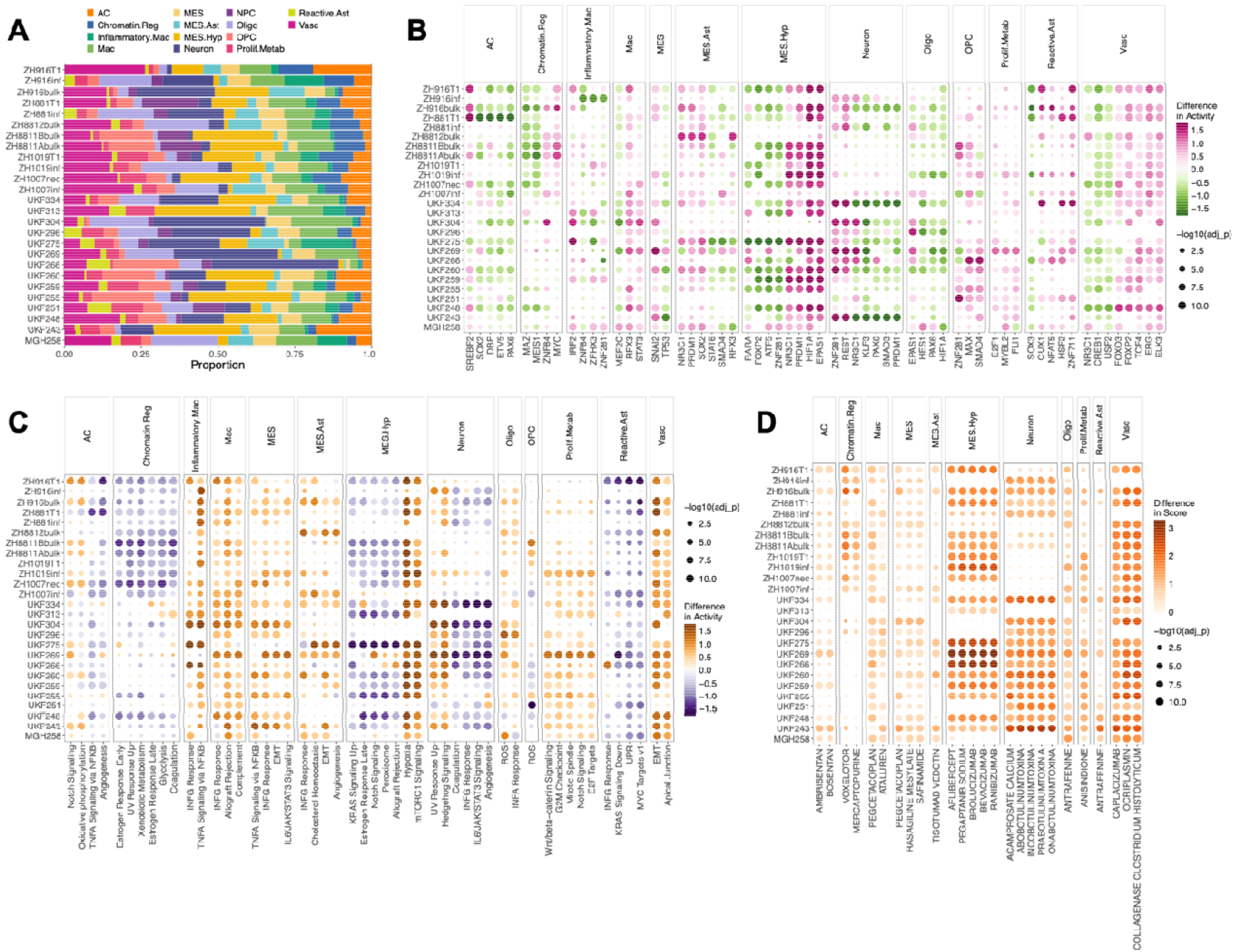
Spatially resolved regulatory programs and therapeutic vulnerabilities across GBM tumors. **(A)** Proportions of spots assigned to malignant and non-malignant metaprograms (MPs) across 26 GBM ST samples. Differential **(B)** TF **(C)** pathway activity across MPs for each sample. Dot color shows the mean activity change of a MP relative to other MPs, and dot size indicates significance (–log10 adjusted *p*). **(D)** Predicted drug response profiles across MPs. Color denotes relative sensitivity and dot size indicates significance. Dot plot of drug scores significantly enriched in specific MP. Significance was determined using a two-sided Wilcoxon test (effect size0□ >□2, *P*□< □0.05, Benjamini–Hochberg corrected).

To uncover the regulatory mechanisms shaping the GBM TME, we imputed spot-specific, spatially coherent TF and pathway activity profiles for each ST sample^10^. We then systematically linked each MP to these inferred regulatory signatures, revealing distinct TF and pathway activity patterns across malignant and non-malignant compartments (Methods, Figure 2B-C). Notably, some MPs are predominantly malignant, while others correspond mainly to non-malignant cell states, enabling direct comparisons of regulatory activity between these compartments. Within MES-Hyp regions, the activity of hypoxia-inducible factors HIF1A and EPAS1 (HIF2A) was significantly higher, indicating a robust transcriptional adaptation to local metabolic stress^12^. This was accompanied by the higher activity of NR3C1 and PRDM1, suggesting a complex integration of stress-response, metabolic, and mesenchymal programs within these unique hypoxic niches. In contrast, SREBF2 (sterol regulatory element-binding protein 2) activity was significantly higher in astrocyte-like regions of the tumor, indicating an active role for lipid metabolism in this cellular state^13^. STAT3^14^ activity was significantly higher in macrophages, consistent with their established roles in regulating macrophage differentiation, inflammatory signaling, and the acquisition of immunosuppressive tumor-associated macrophage phenotypes.

Pathway analysis similarly aligned signaling programs with distinct MPs. MES-Hyp regions showed higher activity of hypoxia and mTORC1 signaling, highlighting metabolic adaptation to hypoxic niches. Inflammatory macrophages showed higher activity of Tumor Necrosis Factor alpha (TNFA)–Nuclear Factor kappa B (NFκB) and Interferon-gamma (IFNG) responses, consistent with immune-activated phenotypes. MES, MES-Ast, and vascular regions showed higher EMT (epithelial-mesenchymal transition) activity, whereas proliferative/metabolic states were defined by higher activity of mitotic spindle and E2F target programs, underscoring their proliferative capacity. Together, these analyses define a spatially resolved regulatory atlas of the GBM TME, directly linking MP-defined cellular states to distinct TF and pathway activities.

### Spatially Resolved Therapeutic Vulnerabilities

To uncover potential therapeutic vulnerabilities associated with distinct cellular states in GBM, we applied drug2cell^*11*^, a computational framework that integrates drug–target interactions from ChEMBL with cell type–specific gene expression profiles. By mapping drug targets onto transcriptionally defined spatial niches and computing drug–cell association scores, this approach enables prediction of candidate therapeutics tailored to localized tumor microenvironments. We focused on clinically approved drugs across all Anatomical Therapeutic Chemical (ATC) classes, selecting those with the strongest predicted effect on a given MP relative to others, and retaining associations consistently observed across samples **(Figure 2D)**. Several predicted associations were concordant with established GBM biology. Hypoxia-driven, angiogenic niches enriched for the MES.Hyp program were predicted to be sensitive to anti-VEGF agents, including bevacizumab, aflibercept, and brolucizumab. Bevacizumab, which is FDA-approved for recurrent GBM where it confers progression-free but not overall survival benefit^15^, provides clinical validation of our approach. Other predictions revealed potential novel vulnerabilities. Core mesenchymal (MES) regions were predicted to respond to monoamine oxidase B (MAO-B) inhibitors^16^, including safinamide, and rasagiline mesylate. MAO-B is overexpressed in high-grade gliomas^17^ and contributes to oxidative stress through hydrogen peroxide production, activating NF-κB signaling and promoting therapy resistance, suggesting a redox-targeted strategy for MES tumors. Macrophage-rich areas were associated with immunomodulatory agents including pegcetacoplan (C3 inhibitor).

Additional associations highlighted MP-specific mechanisms that have received little prior attention in GBM. Chromatin-regulatory programs were linked to antimetabolites (mercaptopurine) and voxelotor, a hemoglobin modulator, potentially reflecting localized proliferation, DNA repair activity, or redox regulation. The astrocyte-like (AC) region was uniquely associated with endothelin receptor antagonists (ambrisentan, bosentan), suggesting a role for endothelin signaling in vascular remodeling or mesenchymal transition. Neuron-like zones were linked to acamprosate, possibly reflecting altered neurotransmission in infiltrative regions, while the oligodendrocyte-like program showed sensitivity only to antrafenine, indicating limited pharmacologic coverage. Proliferation- and metabolism-enriched regions were associated with anisindione, a vitamin K antagonist, whose relevance may involve redox modulation or vascular function.

Collectively, these predictions nominate both established and untested pharmacologic strategies in GBM, grounded in the spatial organization of transcriptional states. This integrative framework provides a blueprint for designing regionally targeted therapeutic approaches that account for intratumoral heterogeneity.

### Ligand–Receptor–Pathway–TF–target gene–Drug Associations in the GBM TME

To dissect the complex cell–cell communication networks within the GBM TME, we applied a systematic approach to model the key components of signaling cascades, including ligands, receptors, pathways, TFs, target genes, and associated drugs. This framework integrates spatially resolved gene expression profiles (e.g., ligands and receptors) with inferred pathway and TF activities, as well as predicted drug sensitivities. Many core TFs driving GBM metaprograms, such as OLIG2, SOX2, and POU3F2, are nuclear transcription factors lacking enzymatic domains or ligand-binding pockets, making them difficult to target directly with conventional small molecules or antibodies. To better understand the regulatory landscape controlling these TFs, we focused on inferring both upstream signaling and downstream transcriptional components associated with their activity.

As a case study, we applied our ligand–receptor–pathway–TF–target gene framework to characterize signaling networks associated with HIF1A activity, a central regulator of hypoxia response and tumor adaptation. For each sample, we correlated HIF1A activity with upstream ligands, receptors, pathway activities, TF activities, downstream target genes, predicted drug scores, and metaprogram proportions (**Figure 3A**). The integrated heatmap revealed that HIF1A-high tumors exhibit coordinated activation across multiple signaling layers. Across ligands, HIF1A activity tracked with angiogenic and hypoxia-responsive, including VEGFA (e.g. **Figure 3B-C**), ADM, ANGPTL4, MIF, S100A10. Receptor/adaptor features co-varying with HIF1A in this figure included RACK1 (GNB2L1), and CAV1. At the pathway level, HIF1A aligned with Hypoxia and mTORC1 signaling, glycolysis, and with TFs characteristic of hypoxic– inflammatory programs, including EPAS1 (HIF-2α), and SOX2. Downstream targets reflected a glycolytic/hypoxia module (GAPDH, ENO1, PGK1, LDHA, TPI1, PKM, MT1X, TMSB10, ENO2, EMP3, NNMT) accompanied by negative associations with myelin/oligodendrocyte genes (MBP, CNP, PLP1). Drug-score correlations highlighted anti-angiogenic agents linked to the VEGF axis—bevacizumab (CHEMBL1201583), brolucizumab (CHEMBL3707357), ranibizumab (CHEMBL1201825), aflibercept (CHEMBL1742982), consistent with the invasive, mesenchymal features observed. Finally, metaprogram proportions in HIF1A-high samples were enriched for mesenchymal/hypoxic and proliferative–metabolic states and relatively depleted for oligodendrocyte/OPC programs, in line with the integrated ligand–receptor, pathway, TF, and target-gene patterns.

**Figure 3.**
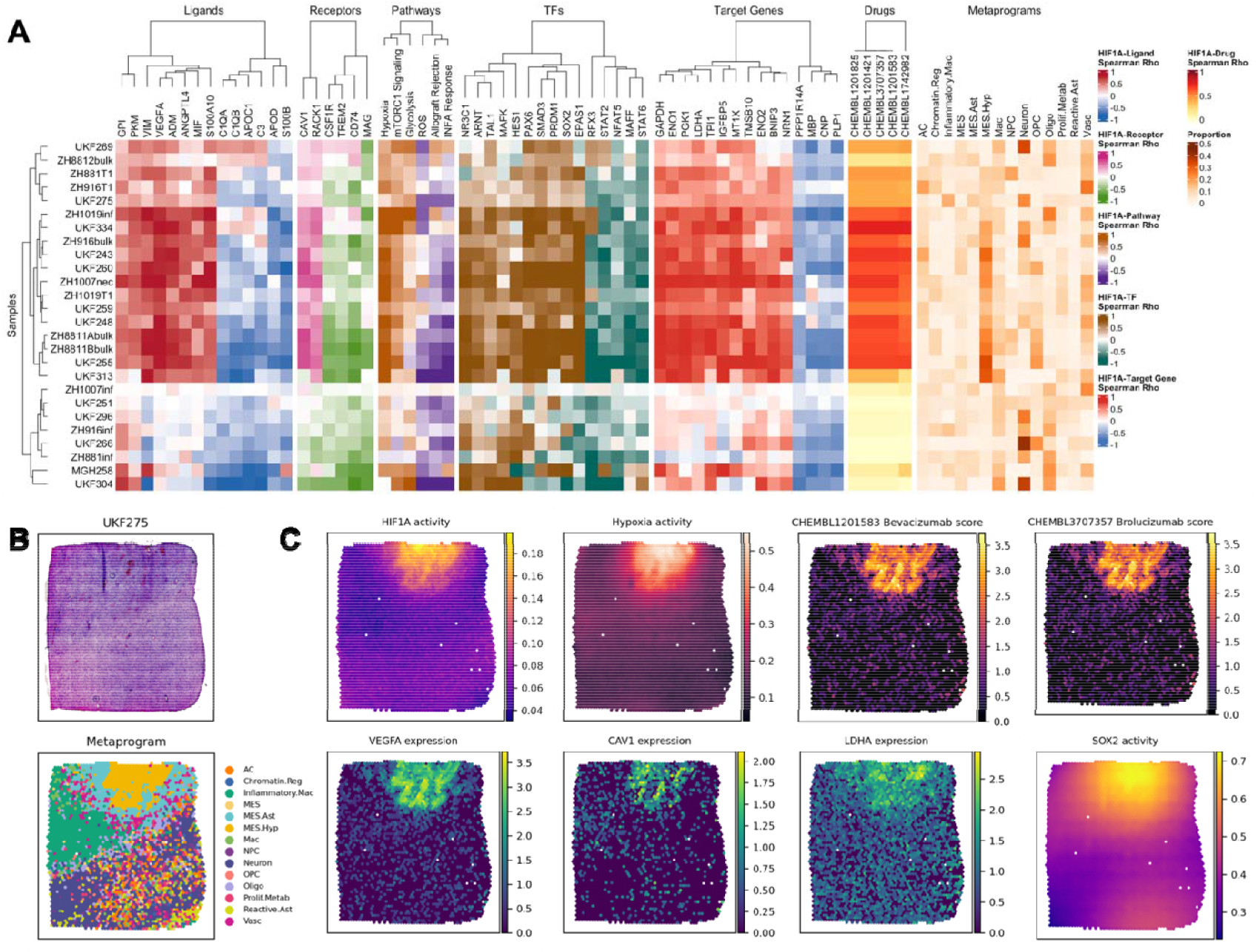
HIF1A-centered multi-layered networks. **(A)** Heatmaps showing sample-wise correlations between inferred HIF1A activity and multiple molecular layers: ligands (L), receptors (R), pathways (P), TFs, target genes (TG), and drugs (D). The rightmost panels depict metaprogram proportions for each sample, providing spatial context for regulator activity. **(B)** Sample H&E-stained image of a GBM sample and metaprogram assignment **(C)** Spatial feature plots displaying the HIF1A, SOX2 TF activity, Hypoxia activity, mRNA expression of VEGFA, CAV1, and LDHA expression and bevacizumab and brolucizumab scores.

Applying the same framework, we also identified a distinct signaling network associated with STAT3 activity. As shown in **Figure 4**, STAT3 activity in glioblastoma exhibited a strong positive correlation with a set of immune-related ligands and receptors. we observed high correlation between STAT3 activity and ligands such as β2-Microglobulin (B2M) and human leukocyte antigens (HLA-A/B), as well as receptors including CD44 and CD74. Pathway enrichment analysis revealed that Interferon Gamma Response, TNF alpha signaling via NF-κB, Epithelial-Mesenchymal Transition (EMT), and IL6-JAK-STAT3 signaling were the most strongly associated pathways, collectively highlighting STAT3’s central role in inflammatory signaling, immune modulation, and mesenchymal transition in glioblastoma. Further investigation into downstream effectors revealed a strong correlation with transcription factors (TFs) like ETV4. These findings provide molecular evidence for STAT3’s established functions in antigen presentation, immune cell recruitment, and immunosuppression. Collectively, these results underscore the multifaceted role of STAT3 as a key orchestrator of the glioblastoma microenvironment, coordinating immune modulation with the promotion of mesenchymal programs.

**Figure 4.**
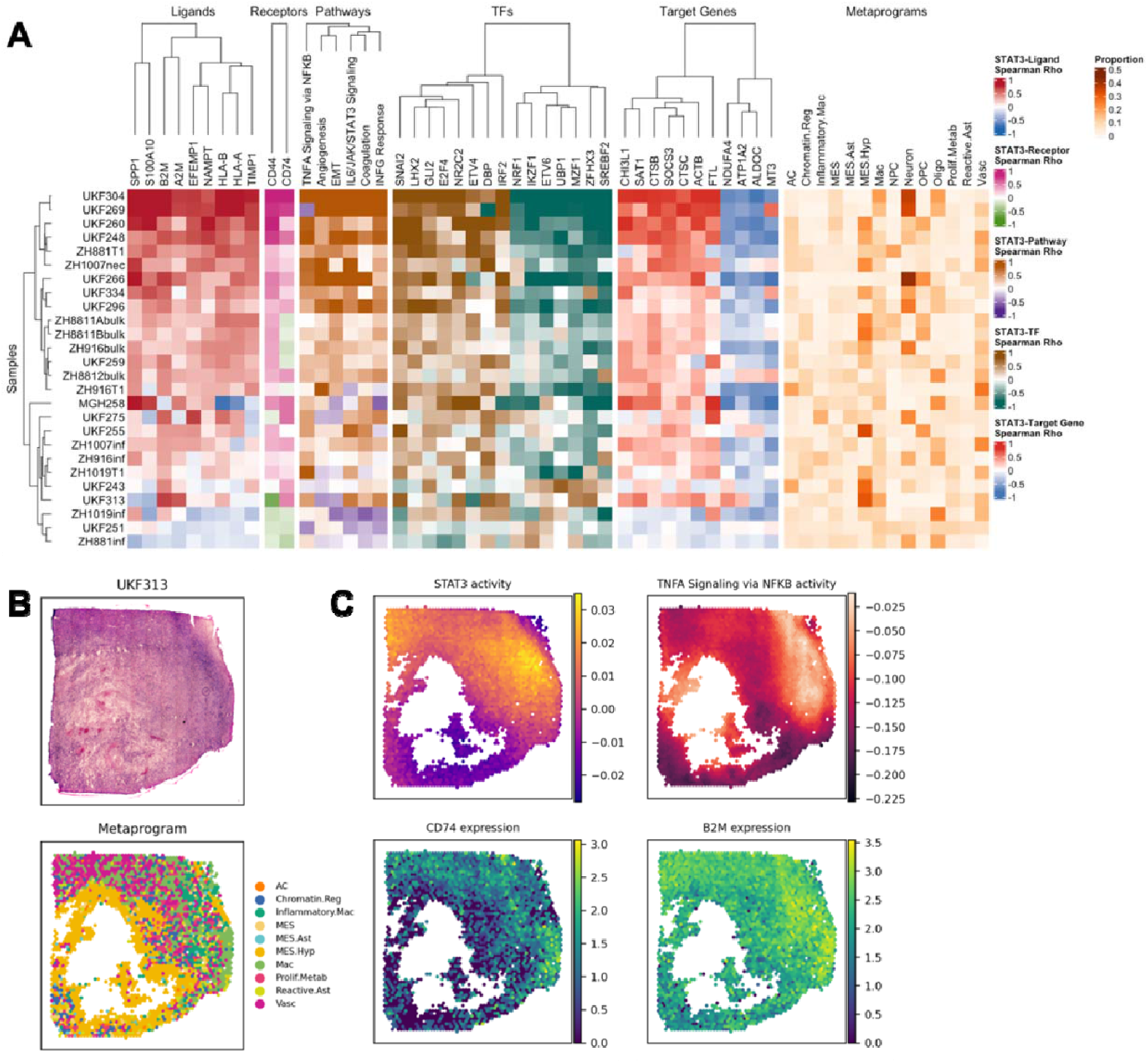
STAT3-centered multi-layered networks. **(A)** Heatmaps showing sample-wise correlations between inferred HIF1A activity and multiple molecular layers: ligands (L), receptors (R), pathways (P), TFs, target genes (TG), and drugs (D). The rightmost panels depict metaprogram proportions for each sample, providing spatial context for regulator activity. **(B)** Sample H&E-stained image of a GBM sample and metaprogram assignment **(C)** Spatial feature plots displaying the STAT3 TF activity, TNFA signaling via NFKB, mRNA expression of CD74 and B2M.

### A spatially resolved, open-source atlas of the glioblastoma tumor microenvironment

To make our analysis accessible for future research, we developed an open, systems-level atlas that empowers the community to test regulatory hypotheses across contexts (https://spatialgbm.streamlit.app/). The platform provides intuitive access to spatial gene expression, metaprograms, TF activity, pathway activity, drug2cell scores and correlation with ligand–receptor expression. Users can explore data by selecting a specific sample or navigating directly to an analysis module from the homepage. This platform empowers researchers to interactively explore the spatial regulation of GBM and supports hypothesis generation for both basic and translational cancer research. Overall, our web resource serves as a comprehensive and accessible resource for the GBM research community.

## Discussion

GBM remains one of the most lethal human cancers, in large part due to the extraordinary heterogeneity of its TME^18^. In this work, we present an integrated, spatially resolved framework that moves beyond expression profiling to construct a multi-layered regulatory atlas of GBM. By integrating STAN^10^ and SPAN with ligand/receptor expression and drug–target predictions^11^, we connect upstream signaling cascades, transcription factor, and pathway activities in a spatially resolved manner. This systems-level integration yields a regulatory roadmap of the GBM TME, linking signaling architecture to transcriptional programs and niche-specific vulnerabilities.

A central insight is the tight coupling of transcriptional metaprograms (MPs) to discrete regulatory signatures. For example, mesenchymal-like (MES) and hypoxia-associated (MES-Hyp) states are coordinated by SOX2^19^, HIF1A, and ARNT together^20^ with pro-tumorigenic pathways such as TNF-α/NF-κB^21^ and mTORC1. These findings support a systems model in which SOX2-high GBM cells adopt a hybrid stem–mesenchymal phenotype that enhances invasion and resistance. HIF1A, in particular, emerged as a master regulator of the hypoxic niche, integrating angiogenic signaling, glycolysis, and mesenchymal transcriptional programs. The predicted sensitivity of HIF1A-high regions to VEGF blockade provides a mechanistic rationale for stratified anti-angiogenic strategies.

Our atlas also nominates additional hubs of therapeutic relevance. Hypoxic MES regions displayed concurrent activation of HIF1A and mTORC1, consistent with stress-adaptive metabolic rewiring. STAT3-high regions exhibited dual associations with inflammatory signaling and immune-modulatory molecules (B2M, HLA-A/B), highlighting STAT3 as a regulator of both immune activation and immune evasion^22^. These findings are particularly salient given the limited efficacy of current GBM treatments. By resolving the regulatory logic of spatial niches, our framework provides a foundation for mechanistically grounded precision strategies, such as targeting the HIF1A–VEGFA–bevacizumab axis or exploiting STAT3-associated immunomodulation.

These findings are particularly salient given the limited efficacy of current GBM treatments. Standard-of-care temozolomide plus radiotherapy yields a median survival of only ∼15 months, and targeted therapies, including bevacizumab, and immune checkpoint blockade, have largely failed to extend outcomes^23^. Intratumoral heterogeneity and cellular plasticity are major contributors to this resistance. By resolving the regulatory logic of spatial niches, our framework provides a foundation for precision strategies such as targeting the HIF1A–VEGFA– bevacizumab axis or exploiting STAT3-associated immunomodulation.

Several limitations warrant acknowledgement. The analysis is based on a modest number of tumors at a single time point, limiting assessment of temporal and inter-patient variation. Visium ST platforms lack single-cell resolution, and therapeutic predictions remain computational. Future work incorporating longitudinal sampling, high-resolution technologies, and functional validation will be essential to translate these predictions into actionable therapies. By making the atlas accessible through an interactive web portal, we enable the community to interrogate multi-layered regulatory programs and nominate testable therapeutic hypotheses. Extensions to cross-species analyses, integration with single-cell atlases, and customizable drug–target inference will further broaden its translational impact.

In summary, our study provides a systems-level lens on the interplay between tumor cells and their microenvironment. Beyond GBM, this atlas establishes a generalizable systems strategy for dissecting spatial regulation across biological contexts. By linking regulatory networks to therapeutic hypotheses in a spatially resolved manner, our framework illustrates how multi-layered maps can guide translational discovery across cancers, immune pathologies, and regenerative contexts.

## Materials and Methods

### Data Acquisition and Pre-processing

We obtained a total of 26 publicly available glioblastoma (GBM) spatial transcriptomics samples from two independent studies^8,9^ profiled on the 10x Genomics Visium platform. The Visium platform profiles gene expression across ∼5,000 spots per section, each 55 µm in diameter and capturing transcripts from ∼1–10 cells. For each of the samples, we acquired the raw gene expression matrix, spatial coordinates, and corresponding histology images. Raw data was processed using Scanpy (v1.11.1)^24^ and anndata (v0.11.4)^25^ in Python (v3.11.5). In brief, we imported the data into Python using Scanpy and anndata and applied a series of quality control filters to remove low-quality spots. Specifically, we filtered out genes detected in fewer than 5 spots and spots with a total count below 1,000. For each filtered gene expression count matrix, we normalized each spot by total counts over all genes in that dataset. The normalized counts were then square root transformed to stabilize the variance.

### Gene Set Curation

For our regulator analysis, we curated a set of well-defined gene sets for both TFs and pathways. For TF analysis, we utilized a comprehensive gene set from the hTFtarget^26^ database, which we further refined to include only TFs listed in the Human Transcription Factor Database^27^. For pathway analysis, we leveraged the Hallmark gene set collection from the Molecular Signatures Database (MSigDB)^28^, providing a robust, biologically coherent source of pathway annotations.

### Spatial Metaprograms

To characterize core transcriptional programs in GBM, we utilized the metaprogram (MP) assignments previously defined by Greenwald et al.^8^. These authors applied non-negative matrix factorization (NMF) to GBM ST samples and identified 14 reproducible MPs, which we adopted without modification. These included eight malignant and six non-malignant programs, reflecting distinct spatial expression patterns. The non-malignant MPs captured canonical cell types and states within the GBM TME: macrophage/microglia (Mac), inflammatory macrophage/neutrophil (Inflammatory-Mac), oligodendrocyte (Oligo), vascular (endothelial/pericyte; Vasc), neuron, and reactive astrocyte (Reactive-Ast). The malignant MPs included two mesenchymal states—MES-hypoxia (MES2) and MES (MES1)—as well as neurodevelopmental-like programs: neural progenitor cell-like (NPC-like), oligodendrocyte progenitor-like (OPC-like), astrocyte-like (AC-like), and a hybrid astrocytic-mesenchymal program (MES-Ast). Additional malignant programs captured proliferative/metabolic activity (Prolif-Metab) and chromatin regulation (Chromatin-Reg).

### Inference of Spatially Resolved Regulatory Activities

We inferred spot-specific TF and pathway activities for each sample using two computational frameworks developed by our group: STAN (Spatial Transcription Factor Activity Network), a pre-existing method^10^, and a new adaptation, SPAN (Spatially informed Pathway Activity Network).

#### STAN for TF Activity Inference

We constructed the TF-target gene prior matrix by configuring the add_gene_tf_matrix function with a minimum gene proportion of 0.1, a minimum of 10 TFs per gene, and a minimum of 5 genes per TF. All subsequent steps were performed using the default parameters.

#### SPAN for Spatially Informed Pathway Activity Inference

To infer spatially informed pathway activities, we adapted the core logic of STAN to develop SPAN. For each sample, the inference process began with a pathway-target gene prior matrix (**P**) (in contrast to the TF-target gene priors used in STAN) and the spot-level gene expression matrix (**Y**). Genes expressed in more than 10% of the spots and present in both matrices were retained. The filtered expression counts were normalized by total spot counts and then square-root transformed to stabilize variance.

For each spot, we constructed a feature vector that combines its spatial coordinates with its local morphological appearance, represented by the mean RGB pixel intensity from a local image crop. These five features were standardized and weighted (spatial:morphological = 1:0.05). We then built a spatial similarity kernel (**K**) using a Gaussian radial basis function (RBF) to calculate the similarity between any two spots based on the distance between their feature vectors.

SPAN assumes that the pathway activity matrix (referred to as **A**) can be split into two matrices: **A = A**_**sd**_ **+ A**_**si**_, where **A**_**sd**_ is spatially and morphologically dependent, and **A**_**si**_ is spatially and morphologically independent. The pathway activity matrix was solved via a spatially weighted regression:

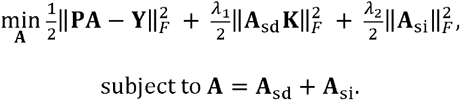

Hyperparameters λ_1_ and λ_2_, were optimized using a custom 10-fold cross-validation grid search on a validation set to achieve the best performance, measured by the Pearson correlation coefficient (PCC) between the predicted and observed gene expression profiles at held-out spots.

### TF–Gene Association Analysis

We correlated inferred STAN TF activities with spatially averaged gene expression using Spearman’s ρ. To focus on robust associations, we retained TF–gene pairs with |ρ| ≥ 0.3 across samples and restricted analyses to genes expressed in at least 5% of spots. Ligands and receptors were annotated from CellTalkDB^2^ to enable downstream TF–ligand and TF–receptor analyses.

### Inference of drug response Associations

We used the drug2cell^11^ Python package (v0.1.2) to identify potential drug–cell type associations. Clinically approved drugs (max_phase = 4) were retrieved from ChEMBL (v.30) and filtered by target organism (*Homo sapiens*), target class, and potency thresholds based on annotations from the NIH Illuminating the Druggable Genome (IDG) project. For each spot, drug scores were computed from the mean expression of their respective target genes. Statistical significance of these scores was assessed using the Wilcoxon rank-sum test, and P-values were adjusted using the Benjamini–Hochberg method. To link our regulatory networks with these drug response predictions, we computed the Spearman’s correlation coefficient (ρ) between our STAN-inferred TF activities and SPAN-inferred pathway activities with the spot-specific drug scores. For subsequent analysis, we filtered out TF/pathway-drug pairs where the absolute Spearman’s correlation was below 0.3 across all samples, to focus on the most robust associations.

### Inferring Metaprogram-specific TFs, pathways, and drugs

To identify metaprogram-specific TFs, we initially scaled the inferred TF activity matrix to unit variance and zero mean. For each metaprogram and each TF, we subsequently conducted a one-sided Wilcoxon rank-sum test to compare the inferred TF activity within and outside the metaprogram. Finally, we adjusted the *P* values via the Benjamini□Hochberg (BH) procedure. The same analytical workflow was applied to the inferred pathway activity matrix and drug2cell score matrix to identify metaprogram-specific pathways and drugs.

For visualization, we selected TFs and pathways for each metaprogram that were both statistically significant (adjusted p-value < 0.001) and differential (difference of activity between the metaprogram and all other spots > 0.3 or < -0.3) in at least 15 samples. From this subset, we further selected 5 TFs with the largest positive mean difference and 5 TFs with the largest negative mean difference in activity. The same procedure and criteria were applied to the drugs, except that we only kept the drugs with positive mean difference in score.

### Statistical Analyses and Visualization

To identify metaprogram-specific TFs, pathways, and drugs, we performed a Wilcoxon rank-sum test for each feature, comparing activity within a metaprogram cluster to all other spots. P-values were adjusted using the Benjamini–Hochberg procedure. To create the TF–gene, TF-ligand, and TF-receptor correlation heatmaps, we computed Spearman’s correlation coefficient for each TF–gene pair per sample. For visualization, we filtered out pairs with an absolute correlation below 0.3 in all samples. We then selected a subset of “molecular features” (ligands, receptors, pathways, TFs, genes, and drugs) based on a high mean absolute correlation (top 20 overall, top 8 for pathways, and top 15 for specific figures) and retained only those with an absolute correlation > 0.5 in at least 5 samples.

All statistical analyses were performed in Python using SciPy^29^ (v1.15.2) and scikit-learn^30^ (v1.5.2) with data manipulation handled by pandas^31^ (v2.2.2) and NumPy^32^ (v1.26.4). Graphs were generated using matplotlib^33^ (v3.10.1), seaborn^34^ (v0.13.2), ggplot2^35^ (v3.5.2), and ComplexHeatmap^36^ (v2.24.1).

### Web Resource Development

We developed a user-friendly web portal to provide interactive access to our analysis results. This open-source resource is freely available at https://spatialgbm.streamlit.app/ without registration. The back end was implemented in Python, while the front end was built with TypeScript (v4.9.3) using the Streamlit (v1.14.0) framework. All charts were generated using in-house Python and R scripts. The portal has been tested for functionality on Google Chrome and Apple Safari browsers. The portal provides a sample-driven interface that allows users to explore the integrated atlas. The homepage features a searchable list of all 26 GBM ST samples. Upon selection, a detailed analysis view displays the H&E histology images, sample metadata, and a navigation panel to access a series of modules. The Spatial Gene Expression module supports visualization of individual gene expression patterns, while the Metaprogram module displays the spatial distribution of the 14 defined malignant and non-malignant transcriptional programs. The TF Activity module allows users to examine spatial patterns of TF activation inferred by STAN, with options to visualize regulators like HIF1A, SOX2, or STAT3. The Pathway Activity module provides access to SPAN-inferred spatial maps of key signaling pathways. The Drug2Cell module matches spatial regulatory profiles to known drug–target relationships. Finally, the Ligand–Receptor-Pathway-TF-target genes-drug association module enables exploration of correlations between receptor–ligand expression and TF/pathway activity. Static figures and logos used in the web portal were created using BioRender (www.biorender.com).

## Data and code availability

Glioblastoma 10X Visium spatial transcriptomics data, including histology images and metaprogram annotations, were downloaded from GitHub (http://github.com/tiroshlab/Spatial_Glioma). A list of human TF-target regulations was obtained from the hTFtarget database^26^ (https://guolab.wchscu.cn/hTFtarget/#!/download). A list of human ligand□receptor interaction pairs was obtained from the CellTalkDB database^37^ (https://github.com/ZJUFanLab/CellTalkDB/blob/master/database/human_lr_pair.rds). The MSigDB Hallmark gene set collection was access through the Python package GSEApy (v1.1.9)^38^. The source codes used to perform the analyses presented in this manuscript are available on GitHub at https://github.com/osmanbeyoglulab/gbm_analysis/.

## Declarations

### Ethics approval and consent to participate

Not applicable (public datasets used).

### Consent for publication

Not applicable.

### Author contributions

HUO designed the study and wrote the paper. LZ performed analyses. ML and XM developed the web platform. All authors contributed to writing and approved the manuscript.

## Acknowledgments

This research was also supported in part by the University of Pittsburgh Center for Research Computing, [RRID:SCR_022735], through the resources provided.

